# Social behavioral deficits in NF1 emerge from peripheral chemosensory neuron dysfunction

**DOI:** 10.1101/790055

**Authors:** Emilia H. Moscato, Christine Dubowy, James A. Walker, Matthew S. Kayser

## Abstract

Neurofibromatosis type 1 (NF1) is a neurodevelopmental disorder commonly associated with social and communicative disabilities. The cellular and circuit mechanisms by which loss of neurofibromin 1 (Nf1) function results in social deficits are unknown. Here, we identify social behavioral dysregulation with loss of Nf1 in *Drosophila*. These deficits map to primary dysfunction of a small group of peripheral sensory neurons, rather than central brain circuits. Specifically, Nf1 regulation of Ras signaling in adult, Ppk23+ chemosensory cells is required for normal social behaviors in flies. Loss of Nf1 results in attenuated ppk23+ neuronal activity in response to pheromonal cues, and circuit-specific manipulation of Nf1 expression or neuronal activity in ppk23+ neurons rescues social deficits. Unexpectedly, this disrupted sensory processing gives rise to persistent changes in behavior lasting beyond the social interaction, indicating a sustained effect of an acute sensory misperception. Together our data identify a specific circuit mechanism through which Nf1 acts to regulate social behaviors, and suggest social deficits in NF1 arise from propagation of sensory misinformation.

## Introduction

Neurofibromatosis type 1 (NF1) is a common and debilitating neurodevelopmental disorder caused by loss-of-function mutations in the neurofibromin 1 gene (*Nf1*) (Xu *et al*., 1990a). *Nf1* is highly expressed in cells of the nervous system, where it functions as a tumor suppressor by negatively regulating the Ras-mitogen activated protein kinase (Ras-MAPK) signaling pathway (Xu *et al*., 1990b; DeClue *et al*., 1992). The disease is predominantly characterized by neurofibromas and other tumors of the nervous system, but broad deficits in neurocognitive functioning significantly degrade quality of life. Compared to the general population, children with NF1 have greatly increased rates of autism spectrum disorder (ASD) (Adviento *et al*., 2014; Plasschaert *et al*., 2015; Morris *et al*., 2016). Studies suggest rates of ASD are 25-50% in NF1 (1-2% in general population), with NF1 patients being 13 times more likely to exhibit highly elevated ASD symptom burden (Morris *et al*., 2016). Social and communicative disabilities stemming from ASDs in NF1 patients are among the greatest contributors to disease morbidity (Plasschaert *et al*., 2015). Children with NF1 experience increased isolation and bullying (Noll *et al*., 2007), difficulties on social tasks, and poorer social outcomes (Barton and North, 2007; Huijbregts *et al*., 2010; Plasschaert *et al*., 2015). Yet, the mechanisms by which loss of *Nf1* results in ASD and social deficits remain largely unknown.

The *Drosophila melanogaster* homologue of *Nf1* is highly conserved, and *Nf1* mutant flies exhibit a range of cellular and behavioral phenotypes with relevance to the human disease. For example, loss of *Nf1* in *Drosophila* is associated with impaired growth, circadian and sleep abnormalities, learning and memory deficits, hyperactivity, and repetitive grooming behavior (The *et al*., 1997; Guo *et al*., 2000; Williams *et al*., 2001; Walker *et al*., 2006; Buchanan and Davis, 2010; Bai and Sehgal, 2015; King *et al*., 2016; van der Voet *et al*., 2016; Bai *et al*., 2018). Disruption of the conserved signaling pathways downstream of *Nf1*, Ras-MAPK and/or cAMP-PKA, has been implicated in these phenotypes. The genetic tractability of *Drosophila* also provides unique opportunities to study how social behaviors are affected by the function of genes, such as *Nf1*, that are associated with neurodevelopmental disorders (NDDs) and ASDs. Previous work has described social deficits with mutation of fly homologs of the Fragile X Mental Retardation gene (*dFMR1*) and ASD-candidate genes such as *neurobeachin* (*rugose*) and *Neuroligin* (*Dnlg-2* and *Dnlg-4*)(Bolduc *et al*., 2010; Hahn *et al*., 2013; Wise *et al*., 2015; Corthals *et al*., 2017). The underlying neuronal circuitry that governs social interactions in flies is precisely mapped (Clowney *et al*., 2015; Kallman, Kim and Scott, 2015), facilitating the examination of how *Nf1* modulates such behaviors with circuit level resolution.

Normally, a male fly responds to a female fly with courtship and to another male fly with rejection. Peripheral sensory neurons in antenna, legs, and mouth detect sex-specific cuticular hydrocarbons (CHCs) on the cuticle of other animals. This flow of sensory information modulates the activity of neurons in the brain to promote or suppress social interactions. In wild-type (WT) *Drosophila melanogaster* males, chemosensory detection of female-specific pheromones activates P1 ‘command’ neurons to promote social interaction and courtship, while detection of male-derived pheromones suppresses P1 activity and inhibits courtship (Billeter *et al*., 2009; Clowney *et al*., 2015; Kallman, Kim and Scott, 2015). Disrupted function of specific groups of sensory neurons leads to aberrant social interactions between flies (Moon *et al*., 2009; Wang *et al*., 2011; Lu *et al*., 2012; Thistle *et al*., 2012; Toda, Zhao and Dickson, 2012; Fan *et al*., 2013; Dweck *et al*., 2015; Hu *et al*., 2015). Interestingly, sensory processing errors and social communication deficits are both prominent symptoms in NDDs including ASDs (American Psychiatric Assocation, 2013), and there is growing recognition that the sensory deficits can actually drive social dysfunction (Hilton *et al*., 2010; Baranek *et al*., 2013; Orefice *et al*., 2016; Ronconi, Molteni and Casartelli, 2016). The cellular and circuit mechanisms that couple sensory and social deficits in NDDs are not well understood.

Here, we use *Drosophila* to examine how *Nf1* functions in a circuit to regulate social interactions. We find that *Nf1* mutant males exhibit social deficits, and that Nf1 acts in a Ras-dependent manner in neurons controlling this behavior. Restoring Nf1 expression only during adulthood rescues social function of *Nf1* mutants, suggesting Nf1 has an ongoing role in coordinating these behaviors in the adult. Behavioral and physiological data reveal Nf1 acts in peripheral sensory neurons, and its loss results in sensory errors underlying social deficits. *In vivo* monitoring of neural activity demonstrates that *Nf1* mutants show decreased chemosensory neuronal activation in response to specific pheromonal cues, giving rise to disinhibition of brain neurons that direct social decisions. Circuit-specific manipulations to restore activity in ppk23+ sensory neurons or suppress activity in P1 courtship “command” neurons both rescue social interaction errors. Surprisingly, disrupted sensory processing in *Nf1* mutants is associated with a persistent behavioral state change that outlasts the social interaction. These findings indicate that social deficits in a fly model of NF1 arise from peripheral sensory neuron dysfunction.

## Results

### *Nf1* mutant males have defects in social interaction behaviors

As an entry point into the function of Nf1 in social functions, we monitored courtship, which consists of stereotyped and selective behavioral routines dependent on social cues. We focused on intermale social interaction by pairing 2 males of the same genotype to determine if mutants are impaired in social behaviors. We first assayed male flies carrying different mutant alleles of *Nf1*. *Nf1^P1^* and *Nf1^P2^* are null alleles generated by mobilization of a P-element (The *et al*., 1997) (Fig 1A). The parental P-element strain, *K33*, served as a control genotype. Mutant flies homozygous for both the *Nf1^P1^* and the *Nf1^P2^* allele displayed a dramatic increase in single-wing extensions, an early step in the courtship ritual in which a male vibrates one wing to produce a mating song, compared to control male pairs which performed few single-wing extensions (Fig 1B; Movie 1-3). Transheterozygous mutants, carrying one *Nf1^P1^* allele and one *Nf1^P2^* allele, also exhibited enhanced single wing extensions. In contrast, heterozygous flies with only one mutated allele of *Nf1* were not significantly different from control flies. We also tested *Nf1^E1^*, an allele generated by EMS-induced mutations in which the protein is truncated upstream of the catalytic domain (Walker *et al*., 2006) (Fig 1A). We assayed transheterozygous flies that carry *Nf1^E1^* and *Nf1^P1^* alleles and found these mutants also show increased single wing extensions (Fig 1B). Together these results demonstrate that loss of Nf1 is associated with aberrant social interaction behavior in male flies.

**Figure 1 (see also Figure S1):**
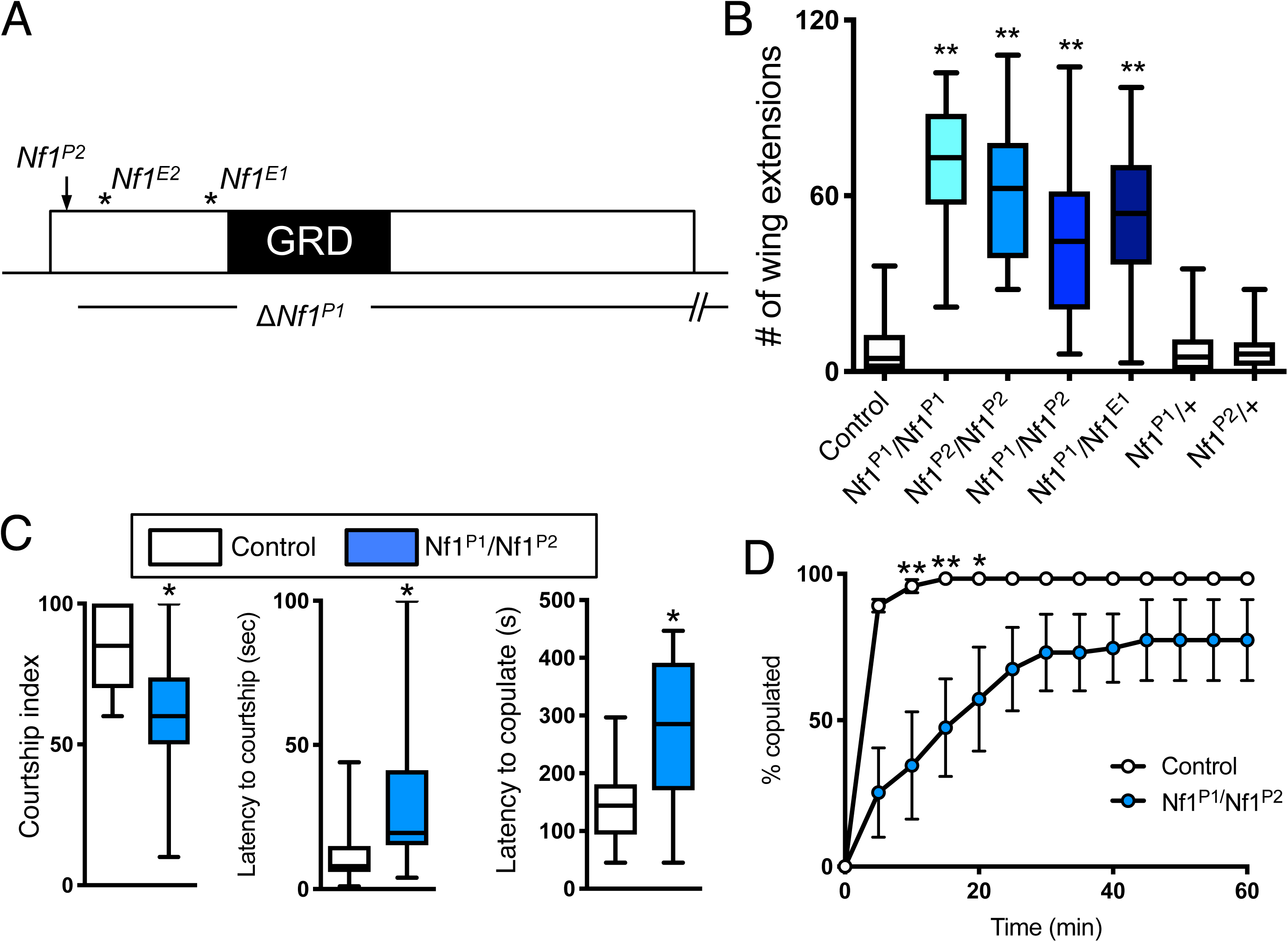
*Nf1* mutant male flies display courtship deficits. A) Schematic of *Nf1* mutant alleles. *Nf1^P1^* has a deletion that removes all of the *Nf1* gene except for the first exon. *Nf1^P2^* contains a P-element in the first intron (arrow). Neither allele expresses Nf1 protein. *Nf1^E1^* and *Nf1^E2^* mutant alleles (asterisks) introduce non-sense mutations upstream of the catalytic domain (GAP-related domain, GRD) and result in a loss of protein product. B) Quantification of single wing extensions in male-male pairs. The genotypes *Nf1^P1^/Nf1^P1^, Nf1^P2^/Nf1^P2^, Nf1^P1^/Nf1^P2^,* and *Nf1^E1^/Nf1^P1^* show a significant increase in the number of wing extensions compared to control flies. *Nf1^P1/+^* and *Nf1^P2/+^* heterozygotes are not significantly different than controls. n = 16-25 pairs per genotype. C) Quantification of courtship parameters in male-female pairs. *Nf1^P1^/Nf1^P2^* males show a decrease in courtship index, an increased latency to begin courting, and an increased latency to copulate compared to control flies. n = 17-20 pairs per genotype. D) Quantification of copulation frequency of control and *Nf1^P1^/Nf1^P2^* mutant male flies paired with females. Mutants are slower to copulate and fewer pairs successfully mate by the end of the assay. n = 45-60 pairs per genotype. *p<0.05, **p<0.01 by Kruskal-Wallis test followed by Dunn’s multiple comparisons test (B), Mann-Whitney test (C), or repeated measures ANOVA followed by Bonferroni’s multiple comparisons test (D).

Do *Nf1* mutant males show enhanced courtship activity toward partners of either sex? Individual *Nf1* mutant males were paired with single WT females in a courtship arena for 10 minutes or until copulation occurred. Courtship index, a measure of the amount of time spent courting a female during the assay, was decreased in *Nf1* mutant male flies compared with control flies (Fig 1C). The latency to initiate courtship was significantly increased in mutant males, as was the latency to copulate (Fig 1C); copulation frequency was reduced (Fig 1D). Thus, *Nf1* mutant males do not simply show increased levels of courtship overall.

We next tested *Nf1* mutant males in competitive copulation assays to determine whether they are less effective than control males at courting a female target. In every assay, the control male ‘won’ over the mutant and copulated with the female (data not shown). We hypothesized that the *Nf1* mutant male flies might ‘lose’ because they are spending time courting the control males, rather than directing their efforts solely toward the female. However, both control and mutant males directed the vast majority of their courtship behavior toward the female target (Fig S1A). This result suggests that *Nf1* mutant male flies are not ‘genderblind’, as has been described in unrelated mutants that also engage in male-male courtship (Grosjean *et al*., 2008). Rather, mutant males can differentiate female from male targets and appropriately direct their courtship efforts toward the female, albeit with less efficacy than control males.

### Loss of Nf1 is associated with impaired chemosensory detection

Altered male-male interactions in pairs of *Nf1* mutants could arise from impaired detection of sensory cues. Alternatively, *Nf1* mutants might produce abnormal cues such that these flies no longer appear male to their partner, leading to nonproductive courtship from other males. For example, male flies without oenocytes fail to produce CHCs and thus elicit courtship from WT males (Billeter *et al*., 2009). To distinguish between these possibilities, we examined whether *Nf1* mutant males elicit single wing extensions from control male tester flies (Fig S1B). If mutants no longer produced the correct blend of CHCs to mark them as male, then we would predict they would elicit courtship from control males. However, we found that control males performed few single wing extensions when paired with either *Nf1* mutants or with other control flies (Fig S1C, left). Mutant flies, in contrast, directed single wing extensions toward both *Nf1* mutants and control flies, although they directed more wing extensions toward other mutant males than control males. We suspected that the apparent preference of *Nf1* mutant males for mutant targets might be due to rejection behavior from control targets. To address this variable, we assessed wing extensions of *Nf1* mutant males paired with decapitated control or mutant male targets (Fig S1C, right). Under these conditions, *Nf1* mutant flies displayed the same level of courtship toward both control and other *Nf1* mutants. These experiments suggest that Nf1 functions in the detection of male cues, rather than the production of those cues within the fly cuticle.

In addition to differentiating between male and female, successful reproduction in the wild requires *Drosophila melanogaster* males to distinguish females of their own species from other drosophilids. Related but distinct species have evolved unique pheromone codes to aid in this recognition: females of the closely related species *Drosophila simulans* produce 7-T, the CHC also produced by *Drosophila melanogaster* males (Jallon and David, 1987; Shirangi *et al*., 2009). 7-T is aversive to *Drosophila melanogaster* males and helps inhibit both intraspecific male-male courtship and interspecific male-female courtship (Lacaille *et al*., 2007; Fan *et al*., 2013). As expected, in a social preference assay, control males preferred to court *D. melanogaster* females over *D. simulans* females (Fig S1D). We found this preference remained intact in *Nf1* mutant males. In contrast, males with a mutation in the chemoreceptor *Gr32a*, previously shown to be necessary for interspecies recognition (Fan *et al*., 2013), showed reduced preference for *D. melanogaster* over *D. simulans* compared to controls. These results suggest that mechanisms for interspecies discrimination are not disrupted in *Nf1* mutant males.

Finally, we tested whether *Nf1* mutant males can appropriately suppress courtship toward a mated female. Recently mated females display traces of aversive male pheromones from their previous copulation that should inhibit potential mates (Tompkins and Hall, 1981; Everaerts *et al*., 2010; Laturney and Billeter, 2016). Over the course of an hour-long pairing with a recently mated female, control males reduced courtship towards the female (courtship index for the last 5 minutes of the assay was significantly decreased compared to the first 5 minutes, Fig S1E). In contrast, *Nf1* mutant male courtship index remained unchanged from the beginning to the end of the assay. Taken together, these data indicate that loss of Nf1 impairs detection of sex-specific cues that should normally serve to direct appropriate social behaviors.

### Nf1 acts in Fru+ neurons to promote normal social interactions

The previous experiments suggest that *Nf1* mutant males cannot detect and/or process inhibitory chemosensory cues from male flies. We next sought to determine where Nf1 is acting. First, we determined whether Nf1 is functioning in neurons. We used a neuron-specific Gal4, *n-synaptobrevin (n-syb)-Gal4*, to re-express Nf1 throughout the nervous system and observed a suppression of *Nf1* mutant male-male single wing extensions (Fig 2A), indicating that Nf1 expression within the nervous system is sufficient to correctly guide social behaviors.

**Figure 2 (see also Figure S2):**
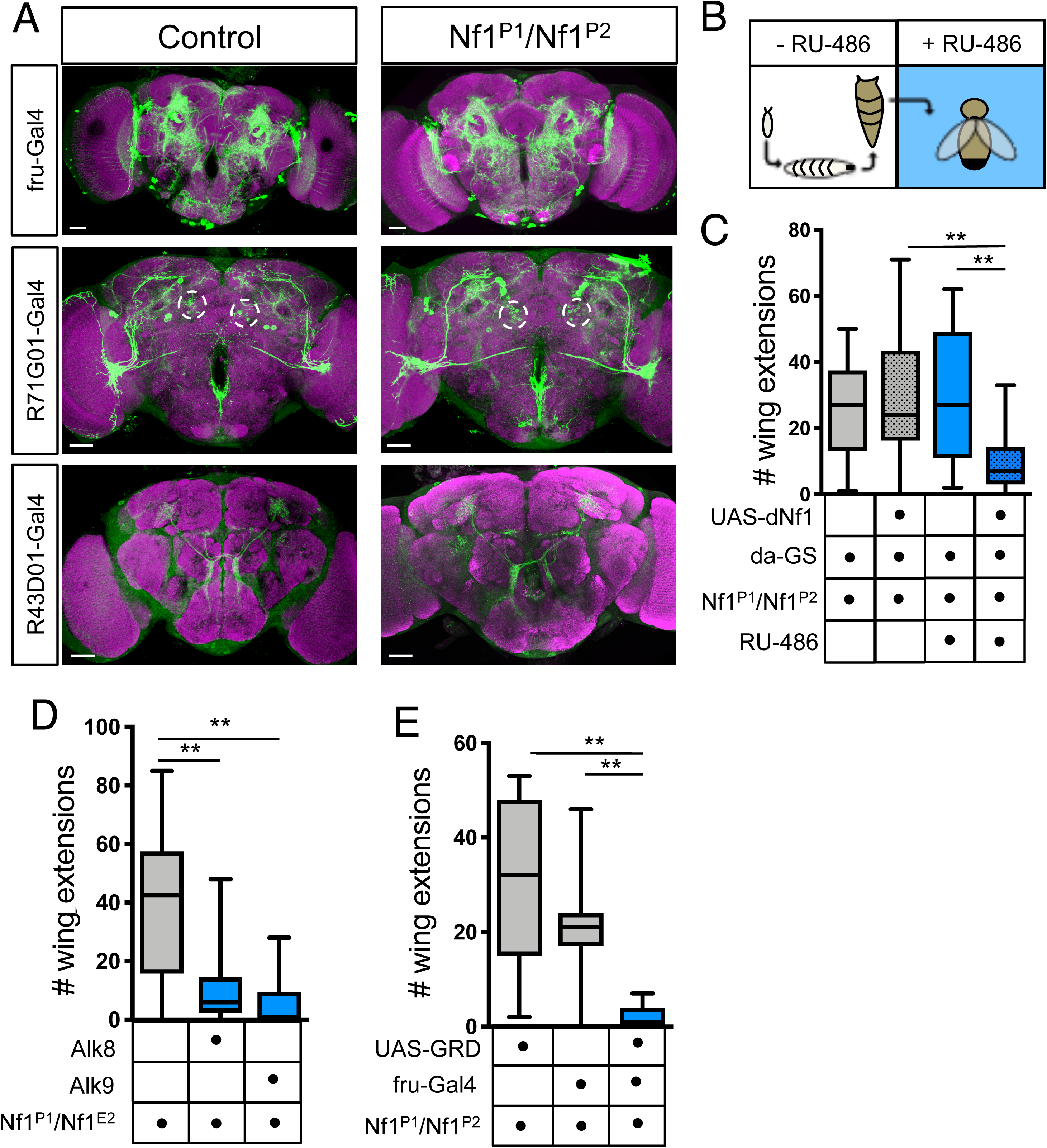
Nf1 acts in Fru+ neurons to promote normal social function A) Quantification of single wing extensions in male-male pairs. When *nsyb-Gal4* was used to drive *UAS-dNf1* pan-neuronally in an *Nf1* mutant background, the number of wing extensions performed by males was significantly decreased compared to *nsyb-Gal4* or *UAS-dNf1* expression alone in a mutant background. n = 13-44 pairs per genotype. B) Quantification of single wing extensions in male-male pairs. When *fru-Gal4* was used to drive *UAS-dNf1* in sexually dimorphic neurons in an *Nf1* mutant background, the number of wing extensions performed by males was significantly decreased compared to *fru-Gal4* or *UAS-dNf1* expression alone in a mutant background. n = 14-21 pairs per genotype. C) Quantification of single-wing extensions in male-male pairs. When *fru-Gal4* was used to knock-down Nf1 expression in sexually dimorphic neurons, the number of wing extensions performed by males was significantly increased compared to *fru-Gal4* or *UAS-Nf1-RNAi* alone. n = 21-22 pairs per genotype. D) When 2 males are paired, detection of volatile and non-volatile pheromones results in activation of mAL neurons, which leads to net inhibition of P1 neurons within the brain. Boxed region shows a schematic of the relevant neural circuitry. E) Quantification of single-wing extensions in male-male pairs. When *R71G01-Gal4* was used to drive *UAS-dNf1* in P1 neurons in an *Nf1* mutant background, the number of wing extensions performed by males was not significantly different compared to *R71G01-Gal4* or *UAS-dNf1* alone in a mutant background. n = 20-28 pairs per genotype. F) Quantification of single-wing extensions in male-male pairs. When *R25E04*- or *R43D01*-*Gal4* was used to drive *UAS-dNf1* in mAL neurons in an *Nf1* mutant background, the number of wing extensions performed by males was not significantly different compared to the *Gal4* or *UAS-dNf1* alone in a mutant background. n = 9-14 pairs per genotype. *p<0.05, **p<0.01 by Kruskal-Wallis test followed by Dunn’s multiple comparisons test (A-C, E-F)

Many of the neural structures and circuits that orchestrate social interaction behavior are sexually dimorphic. A principle transcription factor in the sex-determination hierarchy is *fruitless (fru)* (Demir and Dickson, 2005). The male-specific form of *fru* is expressed throughout the CNS in ∼1700 neurons, including circuits that have been implicated in courtship and other sexually dimorphic social behaviors. We used *fru^NP21^-Gal4* to drive expression of Nf1 in sexually dimorphic neurons in mutant flies, and observed a complete rescue of male-male single wing extensions compared with genetic controls (Fig 2B). To ensure that overexpressing Nf1 in Fru+ circuitry did not broadly suppress all courtship behavior, we tested male-female courtship. Courtship index, latency to copulation, and copulation percentage were indistinguishable between genotypes (Fig S2A-C). Consistent with a role for Nf1 in Fru+ neurons, RNAi-mediated knockdown of Nf1 using *fru^NP21^-Gal4* phenocopied the *Nf1* mutant male social interaction phenotype (Fig 2C). Together, these data show that Nf1 is both necessary and sufficient in sexually dimorphic neural circuits to suppress male-male interactions.

We next sought to more precisely localize Nf1 function within the male central nervous system. The circuitry that transforms socially-relevant sensory information into courtship motor programs is well mapped (Clowney *et al*., 2015; Kallman, Kim and Scott, 2015). Gustatory sensory neurons that detect male pheromones activate mAL interneurons, which in turn inhibit P1 neurons (Fig 2D), a male-specific population of cells known for their ability to trigger courtship behavior. We rescued Nf1 expression in P1 neurons and unexpectedly found no rescue of mutant male-male social interaction behavior (Fig 2E). Expressing Nf1 in mAL interneurons failed to rescue mutant behavior as well (Fig 2F). Thus, restoring Nf1 function to central nodes of courtship circuitry in the brain is not sufficient to normalize *Nf1* male social interaction behavior.

### NF1-related social interaction deficits are caused by disrupted Ras-dependent signaling in adulthood

In humans with NF1, many disease features are thought to be of developmental origin. To distinguish between a developmental versus an adult physiological function of Nf1 in regulating social behaviors, we examined the morphology of courtship neural circuitry in the central brain. We visualized sexually dimorphic neuronal circuitry with GFP (Fig 3A, top panels) and detected no gross changes between controls and mutants. Next we used more restricted expression of GFP to examine specific courtship-related circuits, P1 and mAL, but observed no changes (Fig 3A, middle and bottom panels, respectively).

**Figure 3:**
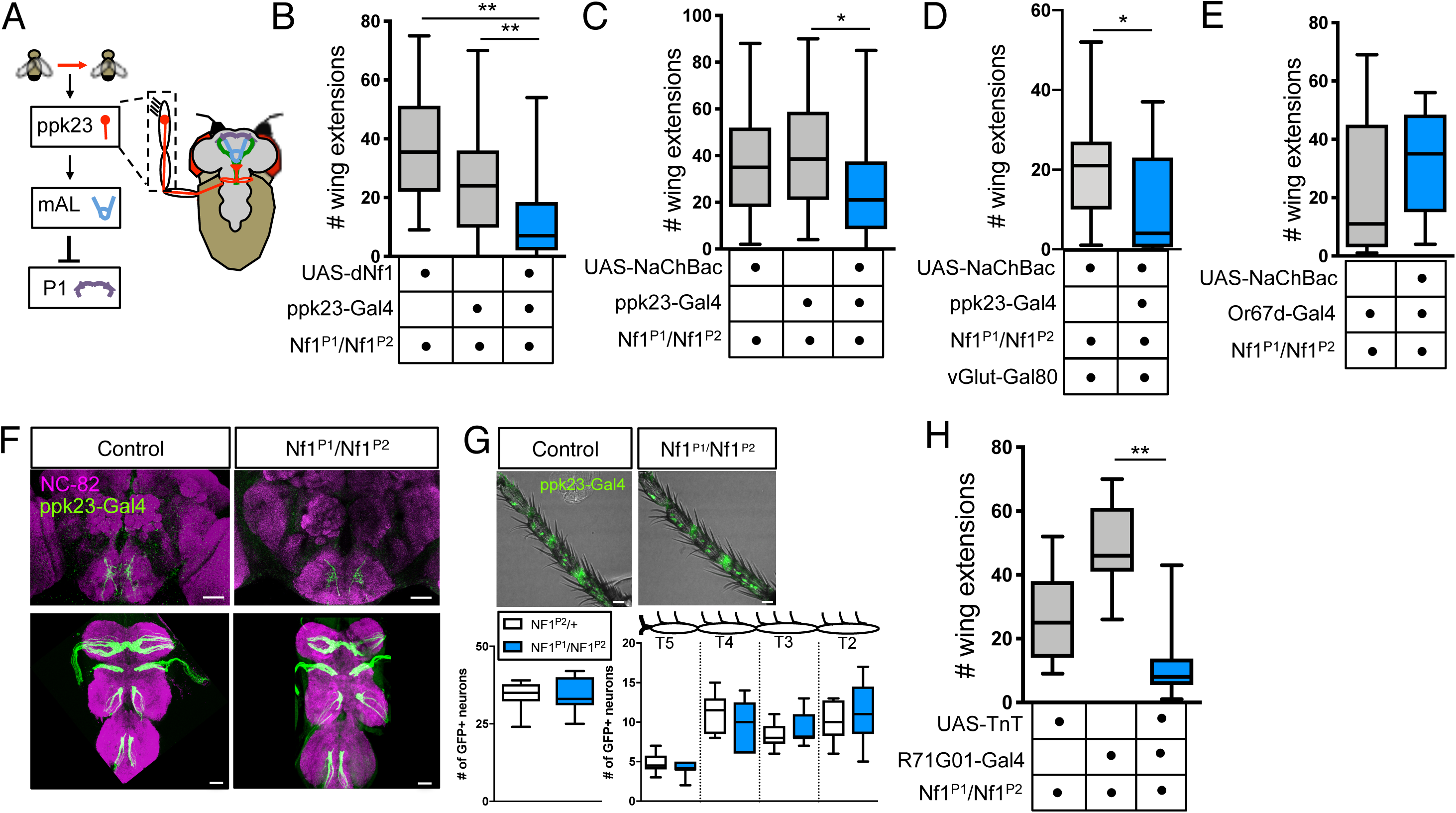
NF1-related social deficits are caused by disrupted Ras-dependent signaling in adulthood A) Courtship-relevant sexually dimorphic circuitry is grossly unchanged in *Nf1* mutant males. Top panels show Fru+ circuitry; left, *fru-Gal4>UAS-mCD8::GFP*; right, *fru-Gal4>UAS-mCD8::GFP; Nf1^P1^/Nf1^P2^*. Middle panels show P1 neurons, relevant cell bodies circled; left, *R71G01-Gal4>UAS-mCD8::GFP*; right, *R71G01-Gal4>UAS-mCD8::GFP; Nf1^P1^/Nf1^P2^*. Bottom panels show mAL neurons; left, *R43D01-Gal4>UAS-mCD8::GFP*; right, *R43D01-Gal4>UAS-mCD8::GFP; Nf1^P1^/Nf1^P2^*. Scale bars 30 μm. B) Schematic of RU-486 time course. Males are not exposed to RU-486 during embryonic, larval, and pupal stages (pre-eclosion), and there is no expression of *UAS-dNf1*. Upon eclosion, adult males are moved to food containing RU-486 to activate gene expression of *UAS-dNf1*. C) Quantification of single-wing extensions in male-male pairs. Grey bars, controls never exposed to RU-486; blue bars, flies exposed to RU-486 post-eclosion. When Gal4-mediated expression of *UAS-dNf1* is turned on by *daughterless-GeneSwitch* (*da-GS*) in a *Nf1* mutant background, the number of wing extensions performed by males is significantly decreased compared to *da-GS* genetic controls (with and without RU-486) and controls of the same genotype not fed RU-486. n = 15-25 pairs per condition. D) Quantification of single-wing extensions in male-male pairs. Loss of one copy of the receptor tyrosine kinase Alk using two different mutant alleles, *Alk8^+/-^* or *Alk9^+/-^*, in an *Nf1* mutant background results in a significant decrease in the number of wing extensions. n = 21-26 pairs per genotype. E) Quantification of single-wing extensions in male-male pairs. When *fru-Gal4* was used to drive just the catalytic domain of Nf1 (*UAS-GRD*) in an *Nf1* mutant background, the number of wing extensions performed by males was significantly decreased compared to *fru-Gal4* or *UAS-GRD* alone in a mutant background. n = 11-30 pairs per genotype. **p<0.01 by Kruskal-Wallis test followed by Dunn’s multiple comparison test (C-E).

Considering that *Nf1* mutant male brains had no obvious morphological abnormalities, we tested if re-expressing Nf1 only in adulthood was sufficient to rescue mutant male social behavior. We used the RU-486 hormone inducible driver *daughterless-*GeneSwitch (da-GS) to gain temporal control over Nf1 expression; in this system, Nf1 is only expressed when flies are fed RU-486. For adult-only rescue of Nf1 function, RU-486 was fed to mutant male flies after eclosion (Fig 3B), which we found was sufficient to restore normal male-male interactions (Fig 3C). Together, our morphological and behavioral data argue against a prominent developmental effect of Nf1 on social behavioral regulation, and instead indicate an on-going role for Nf1 in adult neuronal physiology.

Nf1 operates in a variety of signaling cascades in the central nervous system, most notably in its role as a Ras-GTPase activating protein (Ras-GAP); loss-of-function mutations in Nf1 lead to hyperactive Ras signaling (Martin *et al*., 1990; Xu *et al*., 1990; Bollag *et al*., 1996). In *Drosophila*, mutating the receptor tyrosine kinase Anaplastic lymphoma kinase (dAlk) decreases Ras signaling and rescues size and some behavioral phenotypes in *Nf1* mutant flies (Gouzi *et al*., 2011; Walker *et al*., 2013; Bai and Sehgal, 2015). To test whether elevated Ras signaling contributes to mutant male-male interaction, we assayed *Nf1* mutant male flies that were heterozygous for loss-of-function *dAlk* mutations. Two different mutant alleles of *dAlk* rescued the enhanced single wing extensions of the *Nf1* mutants, demonstrating that normalizing levels of active Ras can reverse social behavioral dysfunction (Fig 3D). To further confirm this finding, we expressed the catalytic domain of Nf1 – the GAP-Related Domain (GRD) – in sexually dimorphic neurons in *Nf1* mutant males, and observed a decrease in single wing extensions (Fig 3E), comparable to the magnitude of rescue with full length Nf1. These data establish that the Ras-GAP activity of Nf1 is sufficient to rescue *Nf1* mutant male social interaction behavior, and that this function can be specifically localized to Fru+ neurons.

### Nf1 is required in Ppk23+ neurons for chemosensory function

Our findings suggest Nf1 acts during adulthood to coordinate social behaviors, but not within examined courtship circuits of the central brain. Disruption of specific sensory neurons in flies results in similar phenotypes to those observed in *Nf1* mutants, so we reasoned that Nf1 might be acting in peripheral sensory neuron function. Our top candidate was Ppk23+ gustatory sensory neurons (GSNs). The *ppk23* gene is a member of the degenerin/epithelial sodium channel (Deg/ENaC) ion channel family, and is expressed in sexually dimorphic Fru+ GSNs (Lu *et al*., 2012; Thistle *et al*., 2012; Toda, Zhao and Dickson, 2012). During social interactions, including courtship, a male fly taps his target using foreleg tarsi to detect CHCs. A subset of tarsal Ppk23+ neurons respond specifically to the male pheromone 7-T, ultimately inhibiting P1 courtship neurons to suppress male-male courtship (schematized in Fig 4A) (Thistle *et al*., 2012; Kallman, Kim and Scott, 2015). Like loss of Nf1, mutations to the *ppk23* gene result in enhanced male-male courtship (Thistle *et al*., 2012; Toda, Zhao and Dickson, 2012). When we rescued expression of Nf1 in Ppk23+ neurons in *Nf1* mutants, we observed a reduction in male-male single wing extensions compared with genetic controls (Fig 4B). This rescue did not reflect a broad reduction in all courtship behavior, as there was no effect on male-female courtship index, latency to copulation, or copulation frequency (Fig S3A-C). As further evidence of a role for Nf1 in Ppk23+ neurons, we tested whether *Nf1* mutant male-male courtship relies on olfactory information. Specifically, aversive CHCs detected by Ppk23+ neurons normally mask attractive olfactory signals and prevent male-male courtship; *ppk23* mutant males lacking antennae show complete suppression of intermale courtship (Thistle *et al*., 2012). Similarly, *Nf1* mutant male-male courtship was nearly abolished in flies lacking antennae (Fig S3D). This finding supports the idea that ectopic courtship in *Nf1* mutant males, as in *ppk23* mutants, results from detection of an attractive olfactory cue that should normally be masked by gustatory detection of aversive CHCs secreted by male flies. Taken together, these data suggest Nf1 normally acts in Ppk23+ neurons to inhibit intermale courtship.

**Figure 4 (see also Figure S3):**
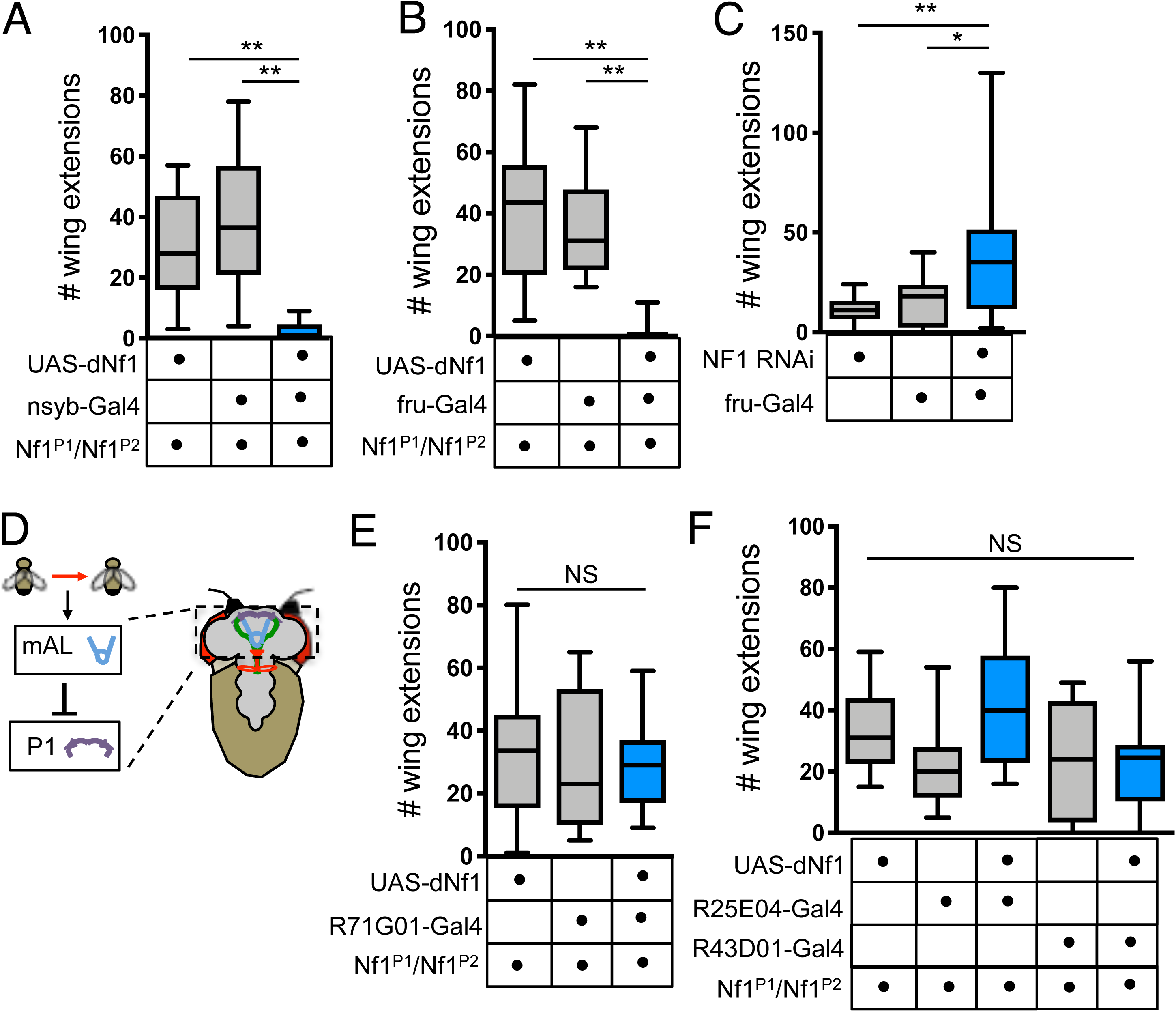
Nf1 functions in Ppk23+ neurons to regulate social behaviors A) When 2 males are paired, detection of non-volatile pheromones results in activation of Ppk23+ gustatory sensory neurons, which leads to activation of mAL neurons and net inhibition of P1 neurons within the brain. Boxed region shows a schematic of the relevant neural circuitry in foreleg tarsi. B) Quantification of single wing extensions in male-male pairs. When *ppk23*-Gal4 was used to drive *UAS-dNf1* in an *Nf1* mutant background, the number of wing extensions performed by males was significantly decreased compared to *ppk23-Gal4* or *UAS-dNf1* expression alone in a mutant background. n = 20-26 pairs per genotype. C) Quantification of single wing extensions in male-male pairs. When Ppk23+ gustatory sensory neurons were constitutively activated with *UAS-NaChBac* in an *Nf1* mutant background, the number of wing extensions performed by males was significantly decreased. n = 25-40 pairs per genotype. D) Quantification of single wing extensions in male-male pairs. When *vGlut-Gal80* was used to restrict expression of *UAS-NaChBac* to Ppk23+ M-cells in an *Nf1* mutant background, the number of wing extensions performed by males was significantly decreased compared to genetic controls lacking *ppk23-Gal4*. n = 17-19 pairs per genotype. E) Quantification of single wing extensions in male-male pairs. When Or67d+ olfactory sensory neurons were constitutively activated with *UAS-NaChBac* in an *Nf1* mutant background, the number of wing extensions performed by males was unchanged compared to *Or67d-Gal4* expression alone in a mutant background. n = 13 pairs per genotype. F) Ppk23+ sensory neuron projections into the ventral nerve cord and brain are grossly unchanged in *Nf1* mutant male flies. Top panels show projections terminating in the sub-esophageal ganglia in the brain, bottom panels show projections through the ventral nerve cord; left, *ppk23-Gal4>UAS-mCD8::GFP*; right, *ppk23-Gal4> UAS-mCD8::GFP; Nf1^P1^/Nf1^P2^*. Scale bars 30 μm. G) Ppk23+ neuronal cell bodies in foreleg tarsi (segments 2-5) in the male fly. Left, *ppk23-Gal4>UAS-mCD8::GFP*; right, *ppk23-Gal4> UAS-mCD8::GFP; Nf1^P1^/Nf1^P2^*. The total number of cell bodies in tarsal segments 2-5 the foreleg tarsi is unchanged between control and mutant males. The number in individual tarsal segments is also unchanged. n = 8-9 flies per genotype, 1 foreleg per fly analyzed. Scale bars 20 μm. H) Quantification of single-wing extensions in male-male pairs. When *R71G01-Gal4* was used to drive *UAS-Tetanus Toxin* (*UAS-TNT*) in P1 neurons in an *Nf1* mutant background, the number of wing extensions performed by males was significantly decreased. n = 7-14 pairs per genotype. *p<0.05, **p<0.01 by Kruskal-Wallis test followed by Dunn’s multiple comparison test (B, C, H); p>0.05, *p<0.05 by Mann-Whitney test (D, E, G).

We hypothesized that in the absence of Nf1, Ppk23+ neurons might not be properly activated in response to male pheromones. In support of this hypothesis, constitutive activation of Ppk23+ neurons with NaChBac, a bacterial sodium channel, resulted in a decrease in *Nf1* mutant male single wing extensions (Fig 4C). The partial rescue could be due to the fact that Ppk23+ neurons comprise two functionally and genetically distinct groups that exert opposing effects on P1 neurons. M-cells respond to male pheromones and inhibit activity of P1 neurons, while F-cells respond to female pheromones and increase P1 activity (Kallman, Kim and Scott, 2015). M-cells can be genetically accessed by combining *ppk23-Gal4* with *vGlut-Gal80*, which suppresses Gal4 expression in F-cells (Kallman, Kim and Scott, 2015). When we selectively activated M-cells in an *Nf1* mutant background, single wing extensions were significantly decreased (Fig 4D). In contrast, activation of Or67d+ neurons, a class of olfactory receptor neurons that also convey an inhibitory signal to P1 upon sensing the volatile male pheromone cVA (Clowney *et al*., 2015), did not rescue *Nf1* mutant male courtship (Fig 4E). These data suggest that in *Nf1* mutant males, Ppk23+ neurons are hypoactive in the presence of male pheromones, and argue that Nf1 functions specifically in peripheral gustatory sensation to modulate social interactions.

We next expressed UAS-mCD8::GFP in Ppk23+ neurons and imaged their axonal projections through the ventral nerve cord into the brain. In *Nf1* mutant males, Ppk23+ axons retained their male-specific midline crossing and terminated in the sub-esophageal ganglion similar to control males (Fig 4F). The number and localization of Ppk23+ cell bodies in the distal 4 foreleg tarsi segments were also identical in control and mutant males as well (Fig 4G). Loss of Nf1 therefore does not disrupt Ppk23+ neuronal morphology.

Ppk23+ neurons are known to provide inhibitory tone to P1 neurons (Clowney *et al*., 2015; Kallman, Kim and Scott, 2015). We wondered whether the *Nf1* mutant behavioral phenotype ultimately emerges from aberrant P1 output due to loss of Nf1-dependent upstream inhibition from Ppk23+ neurons. To test this idea, we examined how experimental inhibition of P1 neurons affects *Nf1* mutant behavior. We found blockade of synaptic output from P1 neurons using Tetanus toxin (TNT) decreased male-male courtship in *Nf1* mutant flies (Fig 4H). These findings suggest P1 hyperactivity contributes to NF1-related social dysfunction, though not due to a role for Nf1 in P1 neurons themselves.

To determine how activity of Ppk23+ neurons following social experience is affected in *Nf1* mutants, we used the CaLexA (Calcium-dependent nuclear import of LexA) system, which drives expression of a GFP reporter in response to sustained Ca^2+^ increases (Masuyama *et al*., 2012). Previously isolated control or *Nf1* mutant males were placed in a vial with wild-type flies of both sexes for 24 hours, after which foreleg tarsi were imaged (Fig 5A). We found that the average number of GFP+, Ppk23+ neurons per foreleg tarsi was significantly decreased in mutant males, though the intensity of the fluorescent signal was unchanged (Fig 5B). These data suggest that during social interactions, Ppk23+ neurons in *Nf1* mutant males are not maximally recruited.

**Figure 5:**
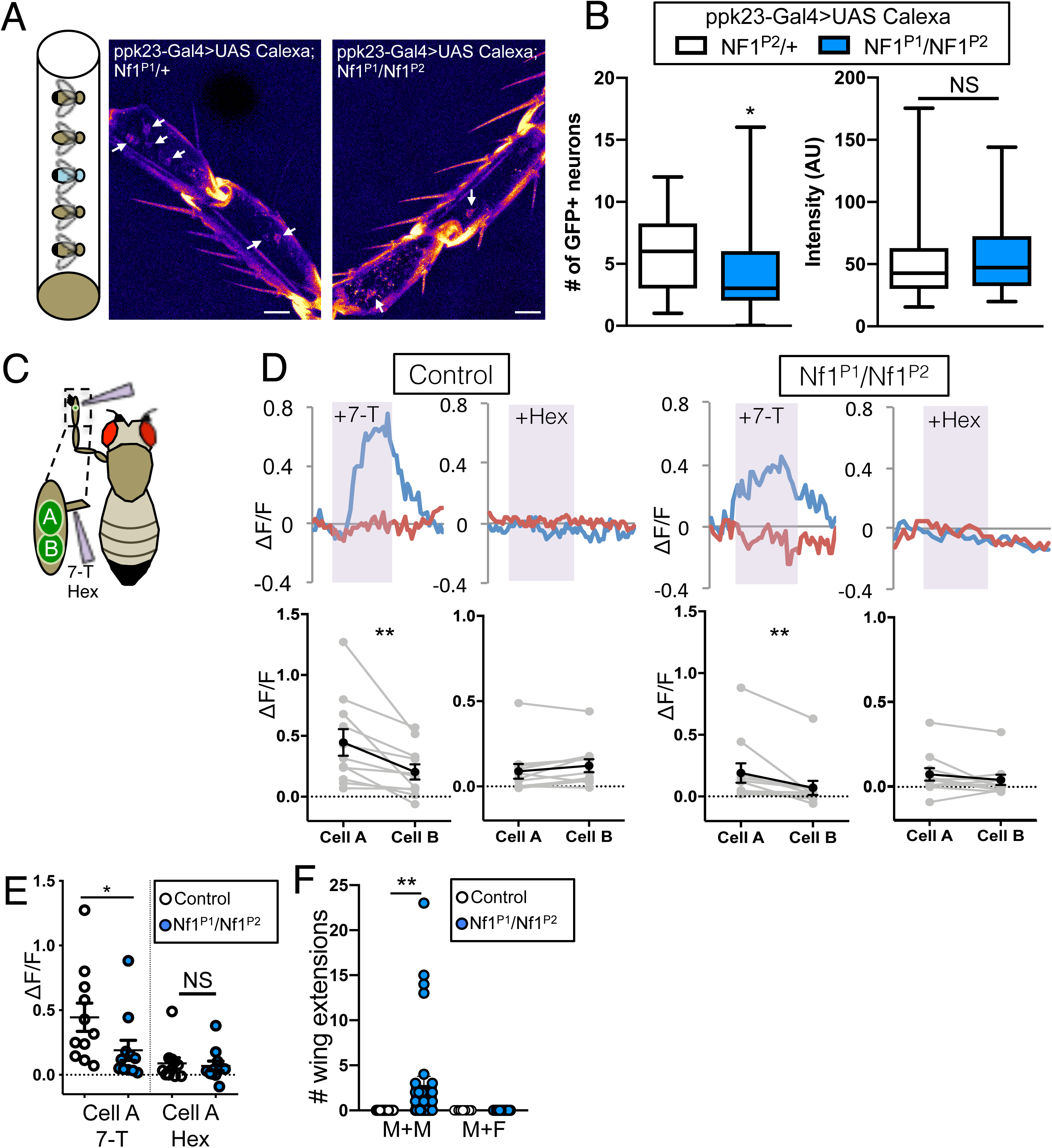
Diminished Ppk23+ activity in *Nf1* mutants in response to social cues A) *Nf1* mutant male flies (blue) expressing *UAS-CaLexA* driven by *ppk23-Gal4* were grouped with wild-type male and female flies for 24 hours (left). Representative images of control and mutant foreleg tarsal segments 2-4, middle and right. Arrows indicate cell bodies with GFP expression. Scale bars 20 μm. B) Quantification of the number (left) and intensity (right) of GFP+ cell bodies in foreleg tarsal segments 2-5. In *Nf1* mutant male foreleg tarsi, the number of activated, GFP+ neurons after 24 hours of social experience is significantly decreased compared to controls. The intensity of the GFP signal is unchanged. n = 27-30 flies per genotype, 1 foreleg per fly. C) Schematic of the experimental preparation for stimulation and imaging of Ppk23+ neurons in foreleg tarsi. A pipet was used to apply 7-T or hexane vehicle to single chemosensory bristles, and *UAS-GCaMP6S* signal in the pair of Ppk23+ neurons underneath was imaged. D) Representative traces (top) and quantification (bottom) of Ppk23+ neuronal responses to 7-T or hexane vehicle in control (left) or mutant (right) male flies. In control flies, upon stimulation with 7-T, there is a large change in the fluorescent GCaMP6S signal in cell A (blue, the putative M-cell) and a significantly smaller response in cell B (red, the putative F-cell). In response to hexane, there is no significant difference in ΔF/F between the two neurons. In *Nf1* mutant flies, there is also a significant difference in GCaMP6s signal between cell A and cell B in response to 7-T, and no change in response to hexane. n = 11 flies per genotype, 1 bristle per fly. E) Quantification of ΔF/F in cell A in control and *Nf1* mutant flies. Control males show a significantly greater response to 7-T than mutant males. There is no change in response to hexane. F) Quantification of single wing extensions after termination of courtship assay by removing the target fly. After a male target is removed from the courtship arena, control flies never perform single wing extensions, while *Nf1* mutant males display a significantly greater number (left, M+M; n = 15-17 per genotype). After a decapitated female target is removed from the courtship arena, flies do not perform single wing extensions (right, M+F; n = 12 per genotype). *p<0.05, **p<0.01 by Mann-Whitney test (B, E, F) and Wilcoxon matched-pairs signed rank test (D).

To directly test the hypothesis that Nf1 is required for detection of male pheromones by Ppk23+ neurons, we monitored calcium signals with GCaMP6s. In males, 2 Ppk23+ neurons, one M-cell and one F-cell, are housed beneath each chemosensory bristle on the foreleg tarsi (Thistle *et al*., 2012; Kallman, Kim and Scott, 2015). Single chemosensory bristles were stimulated with 7-T or hexane vehicle, and fluorescent changes in the Ppk23+ neurons underneath the bristle were analyzed (Fig 5C). As shown previously (Thistle *et al*., 2012), in control males, the calcium increase upon stimulation with 7-T was significantly greater in the male-responsive cell (Cell A) than the female-responsive cell (Cell B) (Fig 5D). The responses to hexane alone were not significantly different between Cell A and B. Similarly, in *Nf1* mutant males we observed a greater response to 7-T in Cell A versus Cell B, and no change in the response to hexane. However, the magnitude of the calcium response to 7-T in the male-responsive Ppk23+ neuron was significantly reduced in *Nf1* mutant males, confirming that Ppk23+ neurons are not properly activated in response to 7-T upon loss of Nf1 (Fig 5D,E). Thus, Nf1 is required for detection of aversive male CHCs by a class of chemosensory neurons, demonstrating a role for Nf1 in gating the flow of sensory input to produce correct behavioral output.

### Persistent behavioral state change in *Nf1* mutants

After initiation of male-male social interactions, we noticed that *Nf1* mutant males frequently continued single wing extensions even when not oriented to the target. Importantly, *Nf1* mutants never exhibited wing extensions in isolation without preceding exposure to a target male. We hypothesized that aberrant sensory function might lead to a persistent behavioral state change. To test this, we paired male flies and allowed them to interact for 5 minutes. We then removed one of the males and monitored the remaining male for 5 more minutes. *Nf1* mutants frequently displayed single wing extensions even after being isolated (Figure 5F; Movie 4-6). In contrast, control males never performed single wing extensions following removal of the target, even when analysis was restricted to flies that performed at least one single wing extension while paired. The absence of sustained wing extensions in control males did not simply reflect low courtship at baseline, as pairing control males with females for 5 minutes (which induced robust courtship) still did not elicit persistent wing extensions (Figure 5F). Surprisingly, although *Nf1* mutant males court a female target, they exhibited no persistent behaviors once the female was removed. These data suggest that during social interactions, improper coding of chemosensory information at the periphery promotes an altered internal behavioral state in *Nf1* mutant males that outlasts the initial error in sensory detection.

## Discussion

Social behavioral deficits represent a common comorbidity in children with NF1, though the etiology remains unknown. Here, we find that *Drosophila Nf1* mutants exhibit social behavioral dysfunction that can be traced to a previously unappreciated cell-autonomous role for Nf1 in chemosensory neurons. Nf1 acts in a Ras-dependent manner in sexually dimorphic, Fru+ neural circuits to inhibit male-male courtship, though not within two main nodes of courtship circuitry in the brain (P1 and mAL neurons). Instead, our data localize Nf1 function to Ppk23+ GSNs in foreleg tarsi, known to be critical for appropriate social decisions (Lu *et al*., 2012; Thistle *et al*., 2012; Toda, Zhao and Dickson, 2012; Fan *et al*., 2013). We observe no gross structural abnormalities in these neurons, and restoring Nf1 expression or manipulating Ppk23+ neuronal activity only in adulthood is sufficient to rescue mutant male courtship behavior, indicating that Nf1 acts in the adult nervous system to promote normal social interactions. Loss of Nf1 results in attenuated response of Ppk23+ neurons to social cues and subsequent non-productive intermale courtship, likely due to failed propagation of a crucial inhibitory signal to the brain. Surprisingly, this behavior persists beyond the social interaction itself, suggesting that an acute sensory misperception gives rise to a sustained behavioral error.

### Sensory processing and social communication in ASDs

In humans, difficulty processing sensory information across modalities greatly impedes social functioning. For example, face and gaze processing impairments make social gestures like eye contact and attention difficult. Atypical auditory processing can disrupt speech recognition and discrimination of emotionally charged speech. Odor preferences can affect the initiation and maintenance of conversation (Thye *et al*., 2017). While our studies were focused on disease mechanisms in NF1, a near universal feature of ASDs is heightened or impaired sensitivity to sensory stimuli (Baranek *et al*., 2014). Not all patients with NF1 harbor ASD symptomatology, but the incidence and severity of ASD in NF1 is strikingly high (Morris *et al*., 2016). Our work therefore provides important insights into the mechanisms of social interaction behavior and sensory processing deficits that could be relevant to the broader population with ASD.

Numerous lines of evidence suggest that dysfunction in peripheral sensory neurons contributes to core behavioral features across a range of ASD models (Han *et al*., 2016; Orefice *et al*., 2016, 2019; Bhattacherjee *et al*., 2017; Oginsky *et al*., 2017; Dawes *et al*., 2018; Perche *et al*., 2018). One influential model is that sensory impairments give rise to errors in brain development, leading to behavioral deficits in adulthood (Orefice *et al*., 2016, 2019). Social and cognitive deficits can be mapped to primary dysfunction of somatosensory neurons rather than neurons within the brain (Orefice *et al*., 2016). Deletion of *Mecp2*, which causes a syndromic form of autism, solely in peripheral mechanosensory neurons leads to behavioral deficits that recapitulate core symptom domains of ASDs; these social behavioral phenotypes are not seen when *Mecp2* is deleted from mechanosensory neurons in adulthood. More recent work suggests the early postnatal period in mice is the critical window during which mechanosensation influences later behavior (Orefice *et al*., 2019); pharmacological approaches to normalize mechanoreceptor function during this developmental time can improve behavioral deficits, including social impairments in *Mecp2* mutant mice.

Our work likewise identifies errors in sensory coding at the periphery as a mechanism of social deficits, but in contrast, primarily implicates an ongoing role for Nf1 in mature sensory neurons. These possibilities are not mutually exclusive. In one model, normal sensory experiences promote typical brain development, and developmentally-timed sensory disturbances result in aberrant brain wiring and behavioral output (e.g. social and cognitive deficits). In another model, neurodevelopmental disorder-associated genes such as *Nf1* have an ongoing role in sensory function; loss of function causes ongoing sensory processing errors, aberrant encoding of social cues to the brain, and behavioral output deficits, all in the absence of structural brain abnormalities. These distinctions point towards different therapeutic strategies; our data suggest adult correction of sensory function in NF1 patients might abrogate certain social deficits.

### Transformation of sensory errors into behavioral output errors

How is a graded change in sensory neuron responsiveness transformed into a dramatic behavioral change? P1 functions as a courtship command center by virtue of receiving input from diverse sensory modalities (contact, vision, olfaction); the summed balance of excitation and inhibition onto P1 dictates behavioral output, not simply the signal strength of each labeled line (Clowney *et al*., 2015; Kallman, Kim and Scott, 2015). In this way, the quantitatively small attenuation of inhibitory Ppk23+ neuron output in the presence of a male leads to fundamentally altered signal structure reaching P1. Genetic silencing of P1 suppresses *Nf1* mutant male courtship, suggesting these neurons are hyperactive in mutants. Future work will directly examine P1 activity in *Nf1* mutants during social interactions, and test whether rescue of Nf1 in Ppk23+ neurons alone restores net P1 suppression by a male target. Supporting this idea, specific activation of Ppk23+ M-cells suppresses mutant intermale courtship. The fly NF1 model will, moreover, facilitate examination of the neural computations taking place at each successive level of the courtship circuit with loss of Nf1.

It is possible that Nf1 acts in other courtship relevant circuits aside from Ppk23+ neurons, and convergence of these aberrant signals amplifies P1 and courtship dysfunction in response to a male cue. Several lines of evidence support this possibility. First, we do not observe complete suppression of male-male courtship with re-expression of Nf1 in Ppk23+ neurons. This is in contrast to rescue using the broader drivers *nsyb-Gal4* or *fru-Gal4*. This difference could reasonably be attributed to variable driver strength and expression levels of Nf1. However, there might be additional neurons in which Nf1 is necessary to fully suppress male-male courtship. There are several other chemosensory neurons that normally detect aversive pheromones and inhibit nonproductive courtship, and Nf1 may affect signaling from these neurons similar to its effects in Ppk23+ neurons. While rescue of Nf1 expression in mAL and P1 does not suppress intermale courtship, Nf1 may also play additional roles in other Fru+ neurons downstream of gustatory sensation. Future studies will more broadly examine Nf1 function within chemosensory systems.

### Dissociable mechanisms of response to male or female cues in *Nf1* mutants

Surprisingly, restoring Nf1 expression to Fru+ or Ppk23+ neurons did not rescue male-female courtship behaviors that we tested (courtship index, latency to copulation, and copulation rates). This suggests that Nf1 does not act only in Fru+ neurons to promote normal male-female social behaviors, but perhaps has a role in Fru- gustatory circuits or modulatory central brain neurons as well (Fan *et al*., 2013; Zhang, Rogulja and Crickmore, 2016). In *ppk23* mutant males, a decrease in male-female courtship is observed along with an increase in male-male courtship (Lu *et al*., 2012; Thistle *et al*., 2012; Toda, Zhao and Dickson, 2012). Therefore Ppk23+ neurons seemed well-positioned to account for both types of courtship deficits observed in *Nf1* mutants. Yet, consistent with *fru-Gal4* rescue data, Nf1 re-expression in Ppk23+ neurons did not rescue male-female courtship. *ppk25* and *ppk29* are two additional members of the Deg/Enac family of ion channels along with *ppk23*, and Nf1 could function downstream of these receptors. However, Ppk25 and Ppk29 are co-expressed in Ppk23+ neurons and act in a complex with Ppk23 to detect the female aphrodisiac 7,11-HD (Liu *et al*., 2018); re-expression of Nf1 using *ppk23-Gal4* should have restored Nf1 function in Ppk25+/Ppk29+ cells. Importantly, *Nf1* mutant males are able to discriminate the sex of their target to preferentially court females, arguing that there is still a robust detection of attractive stimuli from females. *Nf1* mutant males are also able to perform all steps in the courtship sequence, including copulation, demonstrating that the motor output from the central courtship circuitry is intact. A more likely explanation is the decrease in male-female courtship behavior in *Nf1* mutants is secondary to deficits in global processes such as arousal, motivation, or attention, all of which can be affected in NF1 patients.

### Nf1 function in sensory neurons

What intracellular function does Nf1 serve within ppk23+ sensory neurons? There are many steps between ligand binding and intracellular calcium increases, any of which could potentially be impacted by loss of Nf1 signaling. Given that 7-T induces a response in ppk23+ neurons of *Nf1* mutants, albeit diminished, we conclude that ppk23+ channels must still be expressed in the cell membrane. Our data strongly implicate Nf1-mediated Ras signaling as key to its role in sensory function. In the absence of Nf1, hyperactive Ras might modulate ppk23 receptor expression levels or expression of other receptor complex components; alter excitability through phosphorylation-dependent post-translational modification of channels; or interfere with downstream signaling cascades ultimately affecting intracellular calcium. Indeed, previous work suggests a role for Nf1 in regulating intracellular calcium levels in *Drosophila* neurons and mammalian astrocytes (Bai *et al*., 2018). Furthermore, studies from isolated mouse dorsal root ganglion cells demonstrate changes to N-type calcium currents in *Nf1+/-* sensory neurons compared to controls, likely involving post-translational modification to these channels (Duan *et al*., 2014). Related work reported increased expression levels of specific voltage-dependent sodium channels (Hodgdon, Hingtgen and Nicol, 2012). These results suggest increased excitability and transmitter release in cells from *Nf1+/-* mice, perhaps relevant for enhanced pain sensitivity in NF1 (Créange *et al*., 1999; Wolkenstein *et al*., 2001; Wang *et al*., 2005; Hingtgen, Roy and Clapp, 2006). Together, such findings support the idea that Nf1 modulates chemosensory neuron responsiveness in a Ras-dependent manner, and underscore the need for more detailed examination of sensory processing in humans with NF1.

### Persistent behavioral effects of transient sensory misinformation

In *Drosophila*, transient sensory information can lead to persistent behavioral states that outlast the stimulus. For example, brief optogenetic stimulation of P1 neurons can produce a persistent aggressive state in the absence of continued photoactivation (Hoopfer *et al*., 2015). Likewise, single wing extensions continue after termination of P1 neuron photostimulation (Inagaki *et al*., 2014). We find that *Nf1* mutants exhibit persistent courtship behaviors even after termination of a social encounter with another male; this behavior was never observed after interaction with a female target, or in control tester flies following encounters with male or female targets. It will be interesting to determine whether this persistence arises from sustained alterations in activity of P1 neurons following the improperly encoded social stimulus. *Nf1* mutants vigorously court males and females; the dichotomous persistence behavior of *Nf1* mutants following these social interactions reinforces that nuanced disruptions to sensory processes can manifest as pronounced changes in behavioral output. Social deficits in humans with NF1 and other ASDs could similarly reflect sustained alterations to neural encoding of social cues emanating from primary sensory disturbances.

## Acknowledgments

We thank Dr. Michael Fisher for helpful discussions over the course of the project, and Drs. Kimberly Huber, Michael Hart, and Amita Sehgal for input on the manuscript. This work was supported by a McMorris Autism Early Intervention Initiative Fund Pilot Study Award to E.H.M., a Burroughs Wellcome Career Award for Medical Scientists and NIH grant DP2 NS111996 to M.S.K, as well as the New Program Development Award of the Intellectual and Developmental Disabilities Research Center at CHOP/Penn (NIH/NICHD U54 HD086984), with the L. Morten Morley Fund of The Philadelphia Foundation. We thank Drs. Hubert Amrein, Jean-Christophe Billeter, Barry Dickson, Edward Kravitz, Kristin Scott, and Amita Sehgal for reagents.

## Author Contributions

Conceptualization, E.H.M. and M.S.K; Investigation, E.H.M., C.D., J.A.W.,. Writing—Original Draft, E.H.M. and M.S.K.; Writing—Review & Editing, E.H.M., C.D., J.A.W., M.S.K.; Funding Acquisition, M.S.K.; Supervision, M.S.K.

## Declaration of Interests

The authors declare no competing interests.

## Figure Legends

Data are expressed as box plots or mean ± S.E.M. in all figures.

## Methods

### Fly husbandry

Flies were maintained in bottles on standard food at 25°C on a 12h:12h light:dark cycle. *nsyb-Gal4*, *fru^NP21^-Gal4*, *R71G01-Gal4*, *R25E04-Gal4*, *R43D01-Gal4*, *or67d-Gal4*, *UAS-TNT*, *UAS-mCD8::GFP*, *UAS-NaChBac*, *UAS-GCaMP6S*, CaLexA flies, *daughterless-GeneSwitch,* and NF1 RNAi (BDSC #53322) were obtained from Bloomington stock center and outcrossed into an *iso31* background. *Nf1^E1^, Nf1^E2^, Alk^8^* and *Alk^9^* mutant alleles, *UAS-dNf1* and *UAS-GRD* (Walker *et al*., 2006), (Lorén *et al*., 2003) were previously described. Wild-type Canton-S flies were generously provided by Dr. Edward Kravitz. *K33, Nf1^P1^* and *Nf1^P2^* mutant alleles (The *et al*., 1997) were generously provided by Dr. Amita Sehgal. *Gr32a^-/-^* was generously provided by Dr. Hubert Amrein. *Drosophila simulans* stock was generously provided by Dr. Jean-Christophe Billeter. *ppk23-Gal4/CyO* was generously provided by Dr. Barry Dickson. *ppk23-Gal4/TM6,Sb* was generously provided by Dr. Kristin Scott.

### Courtship assays

Male flies were collected as virgins within 4 hours of eclosion and raised in isolation for 4-7 days. Female CS flies were collected as virgins and maintained in vials in groups of 10-15 for 3-5 days. Paired flies were gently aspirated into a well-lit porcelain mating chamber (25 mm diameter and 10mm depth) covered with a glass slide. Assays were performed at 25°C and 50% humidity, recorded for 10 minutes on a Sony camcorder and scored blind to experimental condition. For male-male assays, the total number of single wing extensions was quantified. For male-female assays, courtship index was calculated as the percentage of time a male was engaged in courtship behavior during a period of 10 minutes or until copulation. Copulation percentage was calculated as percentage of flies in each condition that successfully copulated during the 10 min assay. Preference index was calculated as the difference in time spent courting each target divided by the total time spent courting. For copulation frequency (Fig 1), assays were monitored for 1 hour or until copulation occurred. Copulation events were binned into 5-minute intervals. For experiments with mated females, CS females were paired for 48 hours with CS males, then separated under CO2 anesthesia and used 24 hours later in experiment. Assays were recorded for 1 hour and the first and last 5 minutes were scored. For experiments to monitor persistent courtship behavior, flies were paired for 5 minutes, then one fly was removed by aspiration and remaining fly was monitored for an additional 5 minutes. In the M+F condition females were decapitated to prevent copulation. For antenna-less experiments, antennae were surgically removed with forceps 24 hours prior to experiment. In the mock-surgical condition flies were anesthetized for the same period but antennae were left intact. For interspecies discrimination experiments, *Drosophila melanogaster* and *Drosophila simulans* females were anesthetized and labeled with a dot of acrylic paint on the thorax 24 hours prior to assay. For competitive copulation and courtship elicitation experiments, males of different genotypes were labeled with a dot of acrylic paint on the thorax 24 hours prior to assay. For courtship assays using *daughterless-GeneSwitch*, crosses were set up on standard food. Upon eclosion, males were isolated as virgins and transferred to food containing 500 µM RU-486 for 4-7 days until use in experiment.

### Immunohistochemistry and imaging

Brains were dissected in PBS, fixed in 4% paraformaldehyde for 30 min at room temperature, washed in 0.1% PBS-Tx and incubated with primary antibody at 4°C overnight. Following washes in 0.1% PBS-Tx, brains were incubated with secondary antibody for 2 hours at room temperature, washed and incubated with 70% glycerol for 2 hours at room temperature, and mounted in Vectashield. Primary antibodies included mouse anti-nc82 (1:100, Developmental Studies Hybridoma Bank) and rabbit anti-GFP (1:1000, Molecular Probes). Secondary antibodies included donkey anti-rabbit Alexa488 (1:500, Invitrogen) and donkey anti-mouse Alexa647 (1:500, Invitrogen). Brains were visualized with a TCS SP8 confocal microscope and images processed in NIH FIJI/ImageJ. For visualizing GFP fluorescence in forelegs, tarsi were removed with forceps and mounted in vectashield. Imaging was performed within 20 minutes of removal and mounting. For experiments using CaLexA, control or mutant male flies were paired with 10 iso31 females and 10 iso31 males for 20-24 hours. Foreleg tarsi were removed and imaged as above. For quantification of CaLexA signals, cell body areas were manually selected and mean pixel intensity measured.

### GCaMP6S imaging

Tarsi calcium imaging was performed as previously described with slight modifications (Thistle *et al*., 2012). Flies expressing *UAS-GCaMP6S* and *ppk23-Gal4* were mounted on their side to a glass slide with nail polish, and forelegs extended and fixed with thin ribbons of tape over the 1^st^ and 5^th^ tarsal segments, exposing the 2^nd^-4^th^ segments for imaging. Slides were mounted on a TCS SP8 confocal microscope. 7-tricosene (10 ng/ L, μ Cayman chemicals) or hexane vehicle (Fisher) was applied to single chemosensory bristles for 25 seconds with a pipet pulled from a borosilicate glass capillary tube. The maximum change in fluorescence (ΔF/F) was calculated by dividing the peak intensity change by the average Δ intensity 6 seconds immediately prior to stimulation. Cell A and Cell B were defined according to Thistle et al 2012. All image analysis was performed on manually selected ROIs using FIJI/ImageJ.

### Statistics

Data were plotted and analyzed using GraphPad Prism software. Details of statistical tests, including p values and *n*, are included in figure legends.

<u>Movie 1: Control male-male courtship.</u> Control males interacting in courtship assay exhibit rare single wing extensions.

<u>Movie 2: *Nf1* mutant male-male courtship.</u> *Nf1* mutant males interacting in courtship assay exhibit frequent single wing extensions.

<u>Movie 3: *Nf1* mutant male-male courtship.</u> *Nf1* mutant males interacting in courtship assay exhibit frequent single wing extensions.

<u>Movie 4: Control male persistent courtship.</u> Upon removal of one male from courtship assay, remaining control male does not perform single wing extensions.

<u>Movie 5: *Nf1* mutant male persistent courtship.</u> Following removal of one male from courtship assay, *Nf1* mutant males continue to perform single wing extensions in isolation.

<u>Movie 6: *Nf1* mutant male persistent courtship.</u> Following removal of one male from courtship assay, *Nf1* mutant males continue to perform single wing extensions in isolation.

