## Supplemental data for "Social behavioral deficits in NF1 emerge from peripheral chemosensory neuron dysfunction"

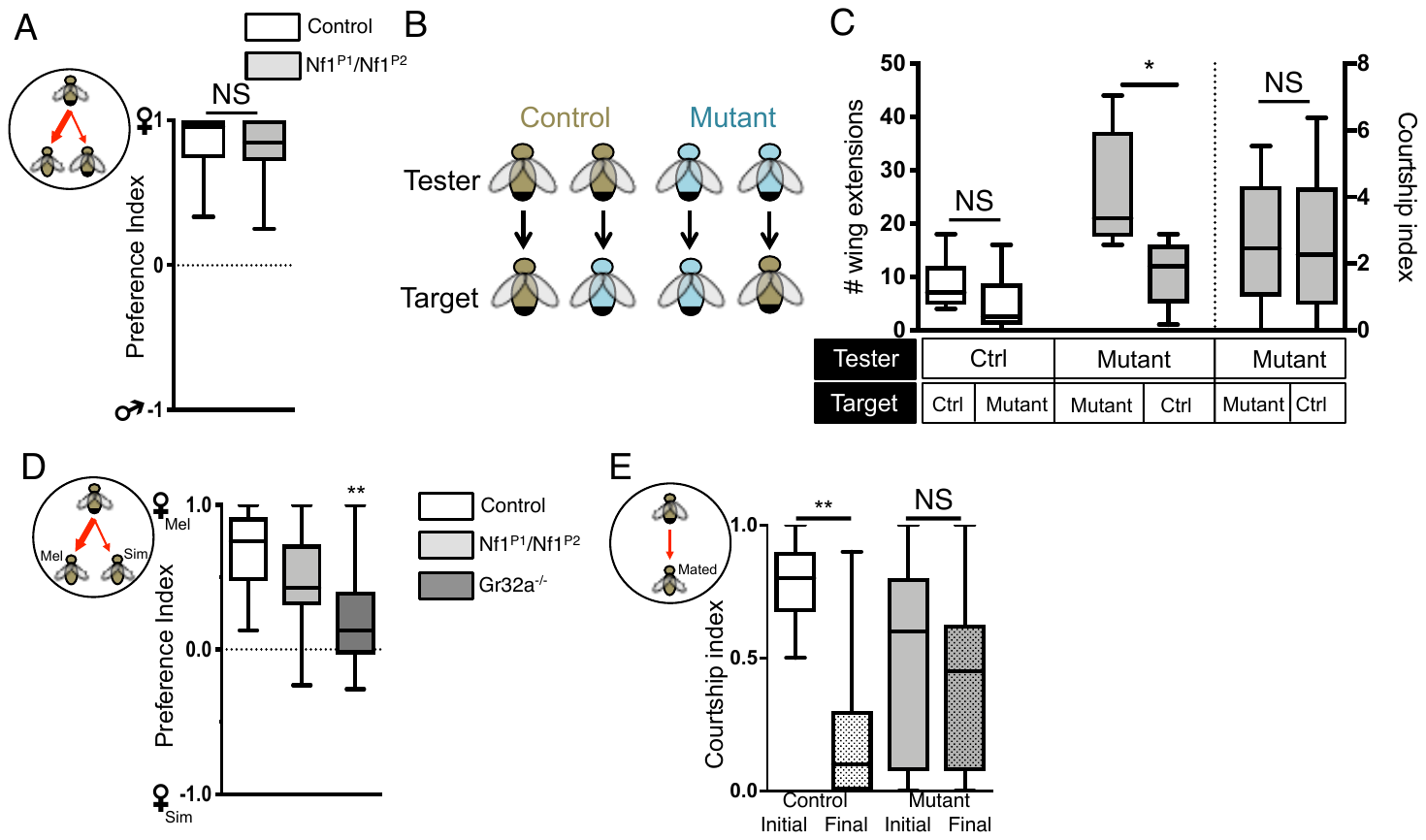

Figure S1. Related to Figure 1: *Nf1* mutant male flies can discriminate sex and species but not mating status.

(A) One control or mutant male was paired with a wild-type male and a wild-type female. Males were distinguish by a dot of acrylic paint on the thorax. Preference index was calculated over a 10-minute courtship assay or until flies copulated and was not significantly different between control and mutant males. n = 17-18 pairs per genotype.
(B) Schematic of experiment. Tester control (brown) or *Nf1* mutant males (blue) were paired with target controls or mutants. Flies were distinguished with a dot of acrylic paint on the thorax.
(C) The number of wing extensions performed by control males toward control or mutant males is not significantly different. Wing extensions performed by mutants toward control males is significantly decreased compared to mutant-mutant pairs. If target flies are decapitated, mutant male courtship index toward a mutant or control target is not significantly different. n = 6-21 pairs per genotype.
(D) One control or mutant male was paired with a *Drosophila melanogaster* female or a *Drosophila simulans* female. Females were distinguished by a dot of acrylic paint on the thorax. Preference index was calculated over a 10-minute courtship assay or until flies copulated. *Nf1^P1^/Nf1^P2^* mutants were not significantly different than controls, while *Gr32a^-/-^* mutants were significantly different. n = 15-26 pairs per genotype.
(E) One control or mutant male was paired with a mated female. Courtship was measured for one hour and courtship index was calculated for the first and last 5 minutes of the assay. Control male courtship index was significantly decreased in the last 5 minutes compared to the first 5 minutes, while *Nf1^P1^/Nf1^P2^* mutant courtship was unchanged. n = 14 pairs per genotype.

**p<0.01 by Mann-Whitney test (A, C), Kruskal-Wallis test followed by Dunn’s multiple comparisons test (D) or Wilcoxon matched-pairs signed rank test (E).

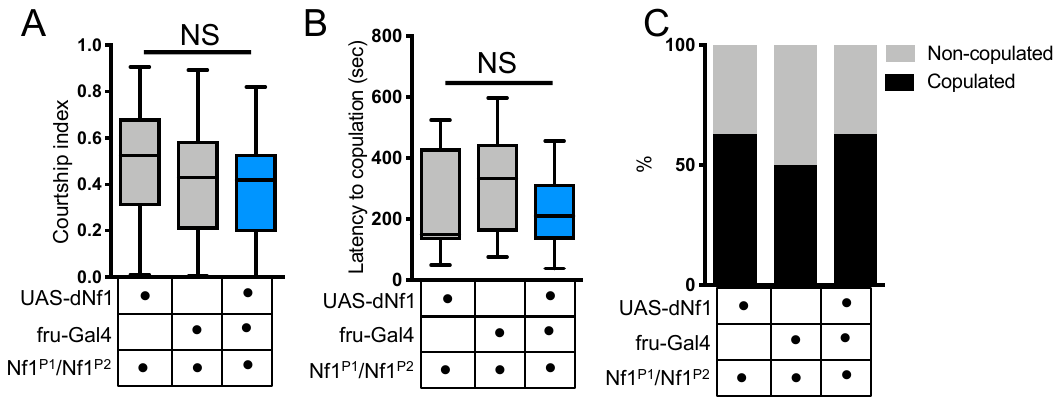

Figure S2. Related to Figure 2: Re-expression of Nf1 in Fru+ neurons does not alter *Nf1* mutant male courtship towards females.

(A) When *fru-Gal4* was used to drive *UAS-dNf1* in an *Nf1* mutant background, courtship index towards a female target was unchanged compared to *fru-Gal4* or *UAS-dNf1* expression alone in a mutant background. n = 26-33 pairs per genotype.
(B) When *fru-Gal4* was used to drive *UAS-dNf1* in an *Nf1* mutant background, copulation latency was unchanged compared to *fru-Gal4* or *UAS-dNf1* expression alone in a mutant background.
(C) When *fru-Gal4* was used to drive *UAS-dNf1* in an *Nf1* mutant background, percentage of male-female pairs that copulated by the end of a 10-minute assay was unchanged compared to *fru-Gal4* or *UAS-dNf1* expression alone in a mutant background.

p>0.05 by Kruskal-Wallis test followed by Dunn’s multiple comparisons test (A, B) or chi-square test (C).

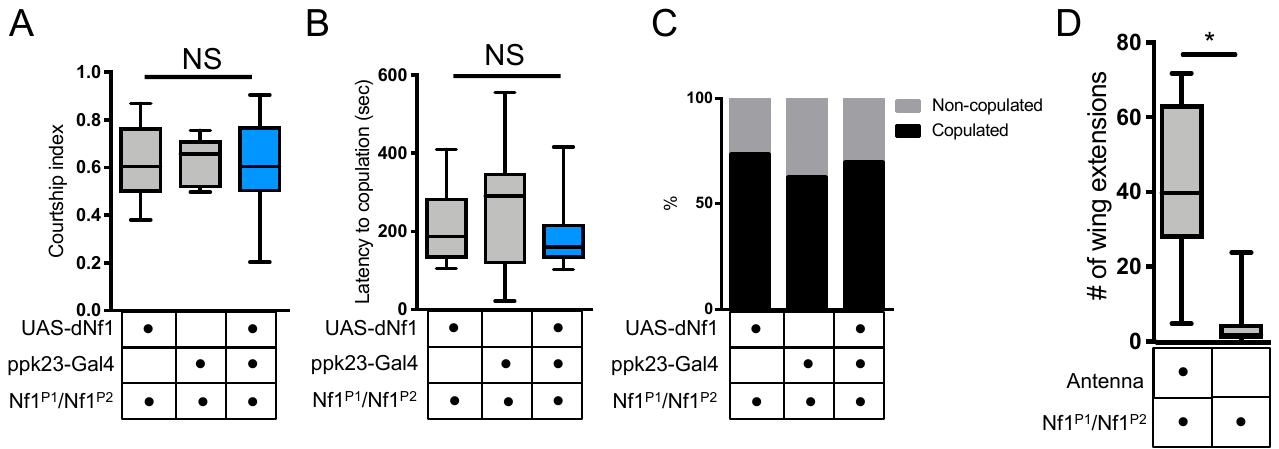

Figure S3. Related to Figure 4: Re-expression of Nf1 in Ppk23+ gustatory sensory neurons does not alter *Nf1* mutant male-female courtship.

(A) When *ppk23-Gal4* was used to drive *UAS-dNf1* in an *Nf1* mutant background, courtship index was unchanged compared to *ppk23-Gal4* or *UAS-dNf1* expression alone in a mutant background. n = 19-25 pairs per genotype.
(B) When *ppk23-Gal*4 was used to drive *UAS-dNf1* in an *Nf1* mutant background, copulation latency was unchanged compared to *ppk23-Gal4* or *UAS-dNf1* expression alone in a mutant background.
(C) When *ppk23-Gal4* was used to drive *UAS-dNf1* in an *Nf1* mutant background, percentage of male-female pairs that copulated by the end of a 10-minute assay was unchanged compared to *ppk23-Gal4* or *UAS-dNf1* expression alone in a mutant background.
(D) Quantification of single wing extensions in male-male pairs. In *Nf1* mutant male pairs, antennectomy causes a significance decrease in courtship index. n = 18-27 pairs per condition.

p>0.05 by Kruskal-Wallis test followed by Dunn’s multiple comparisons test (A, B) or Fisher’s exact test (C); **p<0.01 by Mann-Whitney test (D).
